# Evaluating aerosol and splatter following dental procedures: addressing new challenges for oral healthcare and rehabilitation

**DOI:** 10.1101/2020.06.25.154401

**Authors:** James R Allison, Charlotte C Currie, David C Edwards, Charlotte Bowes, Jamie Coulter, Kimberley Pickering, Ekaterina Kozhevnikova, Justin Durham, Christopher J Nile, Nicholas Jakubovics, Nadia Rostami, Richard Holliday

## Abstract

**Background:** Dental procedures often produce aerosol and splatter which are potentially high risk for spreading pathogens such as SARS-CoV-2. The existing literature is limited.

**Objective(s):** To develop a robust, reliable and valid methodology to evaluate distribution and persistence of dental aerosol and splatter, including the evaluation of clinical procedures.

**Methods:** Fluorescein was introduced into the irrigation reservoirs of a high-speed air-turbine, ultrasonic scaler and 3-in-1 spray and procedures performed on a mannequin in triplicate. Filter papers were placed in the immediate environment. The impact of dental suction and assistant presence were also evaluated. Samples were analysed using photographic image analysis, and spectrofluorometric analysis. Descriptive statistics were calculated and Pearson’s correlation for comparison of analytic methods.

**Results:** All procedures were aerosol and splatter generating. Contamination was highest closest to the source, remaining high to 1-1.5 m. Contamination was detectable at the maximum distance measured (4 m) for high-speed air-turbine with maximum relative fluorescence units (RFU) being: 46,091 at 0.5 m, 3,541 at 1.0 m, and 1,695 at 4 m. There was uneven spatial distribution with highest levels of contamination opposite the operator. Very low levels of contamination (≤0.1% of original) were detected at 30 and 60 minutes post procedure. Suction reduced contamination by 67-75% at 0.5-1.5 m. Mannequin and operator were heavily contaminated. The two analytic methods showed good correlation (*r*=0.930, *n*=244, *p*<0.001).

**Conclusion:** Dental procedures have potential to deposit aerosol and splatter at some distance from the source, being effectively cleared by 30 minutes in our setting.

## Background

The coronavirus disease 2019 (COVID-19) pandemic has had significant impact upon the provision of medical and dental care globally. In the United Kingdom, routine dental treatment was suspended in late March 2020^1–4^, with care instead being provided through a network of urgent dental care centres^5^. During this period, it was advised that aerosol generating procedures (AGPs) were avoided unless absolutely necessary, leading to altered treatment planning and a negative impact on patient care^6^. As more routine dental services start to resume worldwide, the guidance in the UK and elsewhere is still to avoid or defer AGPs where possible^7–13^. This will have an effect both on patients attending for urgent and emergency care, as well as those requiring routine dental treatment for oral rehabilitation. Standard operating procedures (SOPs) have been published by a number of organisations to inform practice, however many of these acknowledge a limited evidence base^14–18^. Additionally, all face-to-face undergraduate and postgraduate clinical dental teaching in the UK is suspended at the time of writing^19^.

Many dental procedures produce both aerosol and splatter contaminated with saliva and/or blood^20, 21^. Saliva has been shown to contain severe acute respiratory syndrome coronavirus 2 (SARS-CoV-2) in infected individuals^22, 23^, many of whom may be asymptomatic^24^, with the salivary gland potentially being an early reservoir of infection^25, 26^. Equally, however, preliminary data suggest that in asymptomatic carriers, the viral load may be low in saliva and these individuals may have faster viral clearance^27, 28^. Early data suggest that SARS-CoV-2 can remain viable and infectious in aerosol for hours, and on surfaces for days^29^. Hence, dental aerosols and splatter are likely to be a high-risk mode of transmission for SARS-CoV-2, and it is highly likely that international clinical protocols across the spectrum of dental practice will need to be significantly modified to allow a safe return to routine care.

A review of the impact of AGPs generally across healthcare (including dentistry) concluded that the existing evidence is limited^30^. The current literature regarding the risks posed by aerosols and splatter in dental settings is particularly limited. A number of authors have used microbiological methods to study bacterial contamination from aerosol and splatter following dental procedures, either by air sampling^20, 31, 32^, swabbing of contaminated surfaces^33, 34^, or most commonly, by collection directly onto culture media^35–38^. These studies are limited in that they only detect culturable bacteria as a marker of aerosol and splatter distribution. A smaller number of studies have used various fluorescent^39–43^ and non-fluorescent tracers^44, 45^ to measure aerosol and splatter distribution, although some of these have significant methodological flaws and major limitations. Many studies are small and report only one repetition of a single procedure, and some have only examined contamination of the operator and assistant; a number of studies which have measured spatial distribution of aerosol and splatter have only done so to a limited distance from the source. Few studies have considered the temporal persistence of aerosol and splatter with sufficient granularity to inform clinical practice.

Open plan clinical environments such as those common in dental (teaching) hospitals with multiple patients and operators in close proximity are problematic. The current lack of robust evidence about dental aerosol and splatter distribution and persistence will be a barrier to the reintroduction of routine dental services and dental education, which is likely to have a negative impact on the availability of care for patients, and on the future dental workforce if not addressed expediently^19^. Patients’ oral healthcare will also suffer if routine care cannot be re-established, especially for those with high dental needs and active dental disease.

The aim of the present study was to establish a robust, reliable and valid methodology to evaluate the distribution and persistence of aerosol and splatter following dental procedures. We present initial data on three dental procedures (high-speed air-turbine, ultrasonic scaler, and 3-in-1 spray use) and examine the effect of dental suction and the presence of an assistant on aerosol and splatter distribution.

## Methods

Experiments were conducted in the Clinical Simulation Unit (CSU) at the School of Dental Sciences, Newcastle University (Newcastle upon Tyne, United Kingdom). This is a 308 m^2^ dental clinical teaching laboratory situated within a large dental teaching hospital. The CSU is supplied by a standard hospital ventilation system with ventilation openings arranged as shown in figure 1; this provides 6.5 air changes per hour and all windows and doors remained closed during experiments. The temperature remained constant at 21.5 °C.

**Figure 1.**
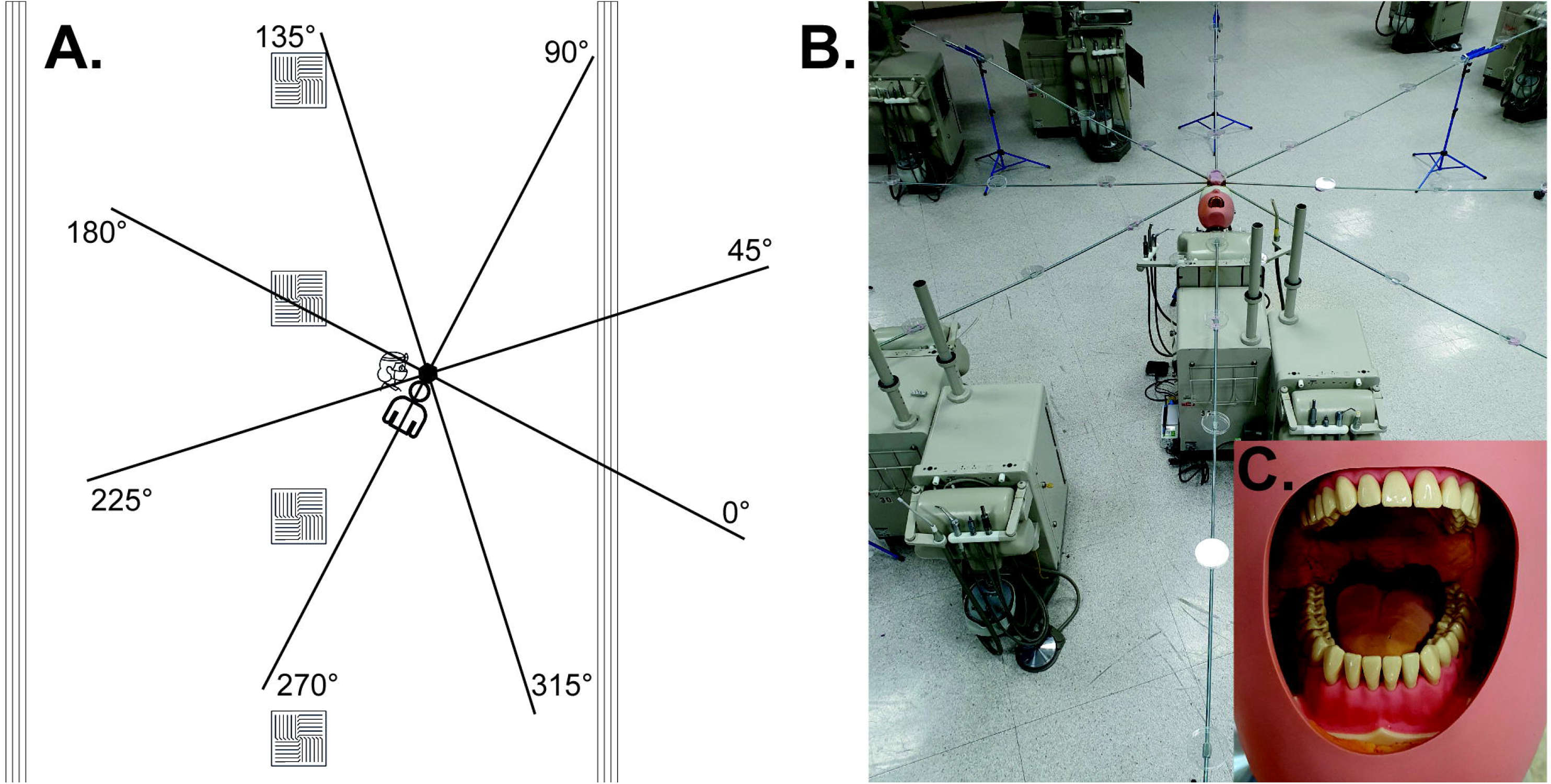
A, Schematic diagram of experimental set up. Position of air vents shown; square vents = air intake; long vents = air output. Experimental set up shown with collection positions labelled (note: degrees are relative to facing the mannequin). B, Photograph of experimental set up showing platforms spaced at 0.5 m intervals to support filter papers. C, demonstration of polyvinyl siloxane addition to mouth of mannequin.

Dental procedures were conducted on a dental simulator unit (Model 4820, A-dec; OR, USA) with a mannequin containing model teeth (Frasaco GmbH; Tettnang, Germany). Polyvinyl siloxane putty (Lab-putty, Coltene/Whaledent; Altstätten, Switzerland) was added to the mouth of the mannequin to recreate the normal dimensions of the oral cavity as described by Dahlke *et al.^41^* (figure 1). Fluorescein solution (2.65 mM) was made by dissolving fluorescein sodium salt (Sigma-Aldrich; MO, USA) in deionised water, and this was then introduced to the irrigation reservoirs of the dental unit and ultrasonic scaler. The procedures investigated were as follows: anterior crown preparation – preparation of the upper right central incisor tooth for a full coverage crown using a high-speed air-turbine (Synea TA-98, W&H (UK) Ltd.; St Albans, UK); full mouth scaling using a magnetostrictive ultrasonic scaler (Cavitron Select SPS with 30K FSI-1000-94 insert, Dentsply Sirona; PA, USA); 3-in-1 spray (air/water syringe) use – washing of mesial-occlusal cavity in upper right first premolar tooth with air and water from 3-in-1 spray. Procedure durations were 10 minutes for anterior crown preparation and ultrasonic scaling, and 30 seconds for the 3-in-1 spray use with air and water (to represent removing acid etchant). Irrigant flow rate was measured at 29.3 mL/min for the air-turbine, 38.6 mL/min for the ultrasonic scaler and 140.6 mL/min for the 3-in-1 spray. We also investigated dental suction (measured at 6.3 L of water per minute) and the presence of an assistant.

Having developed the methods reported by other investigators^36, 41, 43^, the present study used a reproducible, height adjustable rig. This rig was constructed to support cotton-cellulose filter papers spaced at known distances from the mannequin (figure 1). 30 mm diameter grade 1 qualitative filter papers (Whatman; Cytiva, MA, USA) were used to collect aerosol and splatter. These were supported on platforms spaced at 0.5 m intervals along eight, 4 m, rigid rods, laid out at 45° intervals and supported by a central hub, thus creating an 8 m diameter circle around the mannequin; the centre of this circle was located 25 cm superior to the mouth of the mannequin, and in the same horizontal plane as the mouth of the mannequin (73 cm above the floor). Four filter papers were also placed on the body of the mannequin: two at 40 cm from the hub and two at 80 cm. In addition, filter papers were placed on the arms (upper mid-forearm), body (upper chest), and legs (upper mid-thigh) of the operator and assistant as well as on their full-face visor (width: 28.0, height: 27.5 cm) and the vertex of the head. For one condition (anterior crown prep with suction and assistant) we also placed three filter papers on the mask of the operator/assistant (beneath a full-face visor). Two operators conducted the procedures: RH conducted the high-speed air-turbine and ultrasonic scaler procedures (operator height = 170 cm); JRA conducted the 3-in-1 spray procedure (operator height = 175 cm). There was a single assistant with a height of 164 cm.

Before each procedure the mannequin, rig and filter paper platforms were cleaned with 70% ethanol and left to fully air dry. A period of 120 minutes was left between each procedure to allow for clearance of aerosol and splatter. Following each procedure, the filter papers were left in position for 10 minutes to allow for settling and drying of aerosol and splatter, before being collected with clean tweezers and placed into a single-use, sealable polyethylene bag. For the anterior crown preparation without suction, additional filter papers were placed at 30 minutes and again at 60 minutes to examine persistence of aerosol and splatter. At both of these time points, the risk of fluorescein transfer was minimised by placing the new filter papers on new platforms, and filter papers were then left for 10 minutes before collection. All experimental conditions were repeated three times.

### Image Analysis

Filter papers were placed on a glass slide on a black background, covered by a second glass slide, and illuminated by two halogen dental curing lights (QHL75 model 503; Dentsply, NC, USA) with 45 mW/cm^2^ output at 400-500 nm; these were positioned at 0 and 180 degrees, 5 cm from the centre of the sample horizontally, and 9 cm vertically, with both beams of light focussed on the centre of the sample. Images were captured with a digital single-lens reflex camera (EOS 1000D, Canon; Tokyo, Japan) at 90 mm focal length (SP AF 90mm F/2.8 Di Macro, Tamron; Saitama, Japan) with an orange lens filter, positioned 43 cm directly above the sample (sample to sensor). Exposure parameters were f/10, 1/80 seconds and ISO 400. Image analysis was performed using ImageJ^46^ (version 1.53b, U.S. National Institutes of Health; MD, USA) in a darkened room by one of four examiners blind to experimental conditions and sample position (JA, CC, DE, RH). Images were converted into 8-bit images and the pixel scale set across the maximum diameter of the sample at 30mm. A manual threshold was used to create a mask selecting all high intensity areas. The “analyse particles” function was used to identify particles from 0-infinity mm^2^ in area and 0-1 in circularity. The number of particles, total surface area, and average particle size were calculated. Total surface area was selected as the primary outcome measure, representing contamination levels of the samples. Examiners underwent calibration prior to formal analysis by independently analysing 10 images and then discussing to reach consensus. Following this, examiners then independently analysed 30 images to assess inter-examiner agreement. Examiners re-examined the same 30 images one week later to assess intraexaminer agreement.

### Spectrofluorometric Analysis

For one experimental condition (anterior crown preparation without suction, samples from the initial, 30-, and 60-minute time points) we completed spectrofluorimetric analysis to allow validation of the image analysis technique. Building on the methods reported by Steiner et al.^47^, fluorescein was recovered from filter papers by addition of 350 μL deionised water. Immersed samples were shaken for 5 min at 300 rpm using an orbital shaker at room temperature. The fluorescein was then eluted by centrifugation at 15,890 *g* for 3 min using a microcentrifuge. 100 μL of the supernatant was transferred to a black 96-well microtitre plate with a micro-clear bottom (Greiner Bio-One; NC, USA) in triplicate in order to measure fluorescence. Fluorescence measurements were performed using a Synergy HT Microplate Reader (BioTek; VT, USA) at an excitation wavelength of 485 ± 20 nm and an emission wavelength of 528 ± 20 nm with the top optical probe. For background correction, negative controls (n = 26) were included in the measurements for all runs. These included fresh filter papers out of the box and filter papers that had been placed on platforms in CSU for 10 minutes exposed to air. The negative control filter papers were processed for imaging and fluorescent measurements in the same manner as the remainder of samples. The negative control mean + 3SD (164 RFU; relative fluorescence units) was used as the limit of detection; hence a zero reading was assigned to values below 164 RFU. For readings above the detection limit of the instrument (>100,000 RFU), a value of 100,000 RFU was assigned.

### Statistical Methods

Data were collected using Excel (2016, Microsoft; WA, USA) and analysed using SPSS (version24, IBM Corp.; NY, USA) using basic descriptive statistics and Pearson’s correlation (to compare analytical techniques). Heatmaps demonstrating aerosol and splatter distribution were generated using Python 3^48^. A two-way mixed effects model was used to assess inter- and intra-examiner agreement by calculating interclass correlation coefficient (ICC) using STATA release 13 (StataCorp; TX, USA).

## Results

### Examiner Calibration for image analysis

Inter-examiner ICC for 30 images showed excellent agreement for total surface area (ICC 0.98; 95%CI 0.97-0.99), good agreement for total number of particles (ICC 0.88; 95%CI 0.80-0.93), and moderate agreement for average particle size (ICC 0.63; 95%CI 0.47-0.78). Intra-examiner agreement at one week for the same 30 images was excellent for total surface area (ICC 0.97-0.99), good to excellent for total number of particles (ICC 0.82-0.97), and good for average particle size (ICC 0.75-0.97)^49^.

### Aerosol and splatter distribution

Aerosol and/or splatter deposition (assessed by surface area outcome) was highest at the centre of the rig and decreased with increasing distance from the centre (table 1). Most contamination was within 1.5 m but there were smaller readings up to 4 m for some conditions. The spatial distribution is shown in figures 2 and 3.

**Figure 2.**
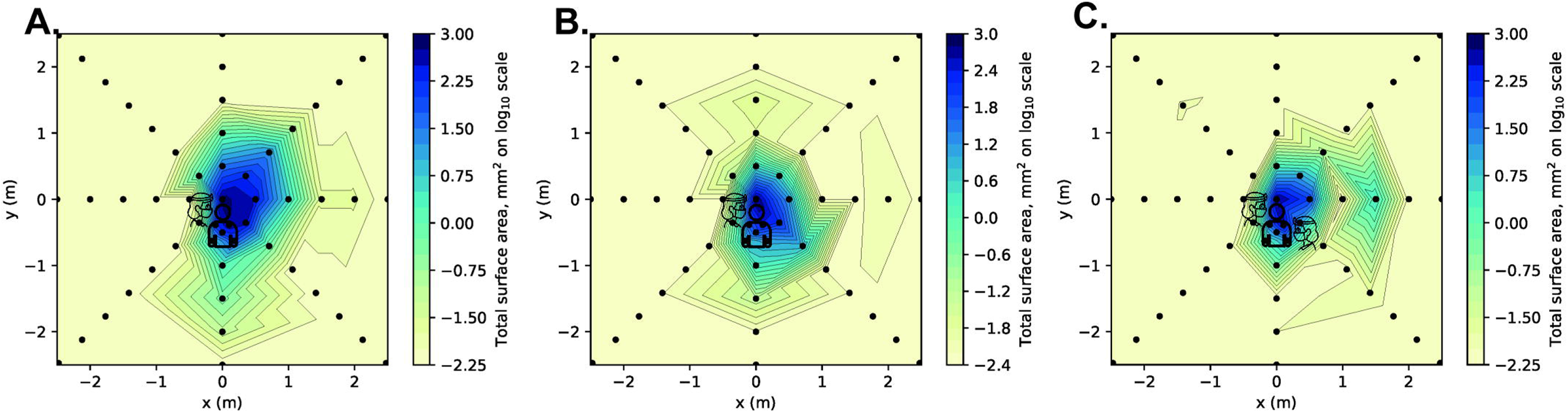
Heatmap showing surface area outcome measure for three clinical procedures. A, anterior crown preparation (without suction). B, anterior crown preparation with suction. C, anterior crown preparation with suction and assistant. For each coordinate, the maximum value recorded from three repetitions of each clinical procedure was used as this was deemed most clinically relevant. Logarithmic transformation was performed on the data (Log_10_). Note the scale is reduced to remove areas showing zero readings.

**Figure 3.**
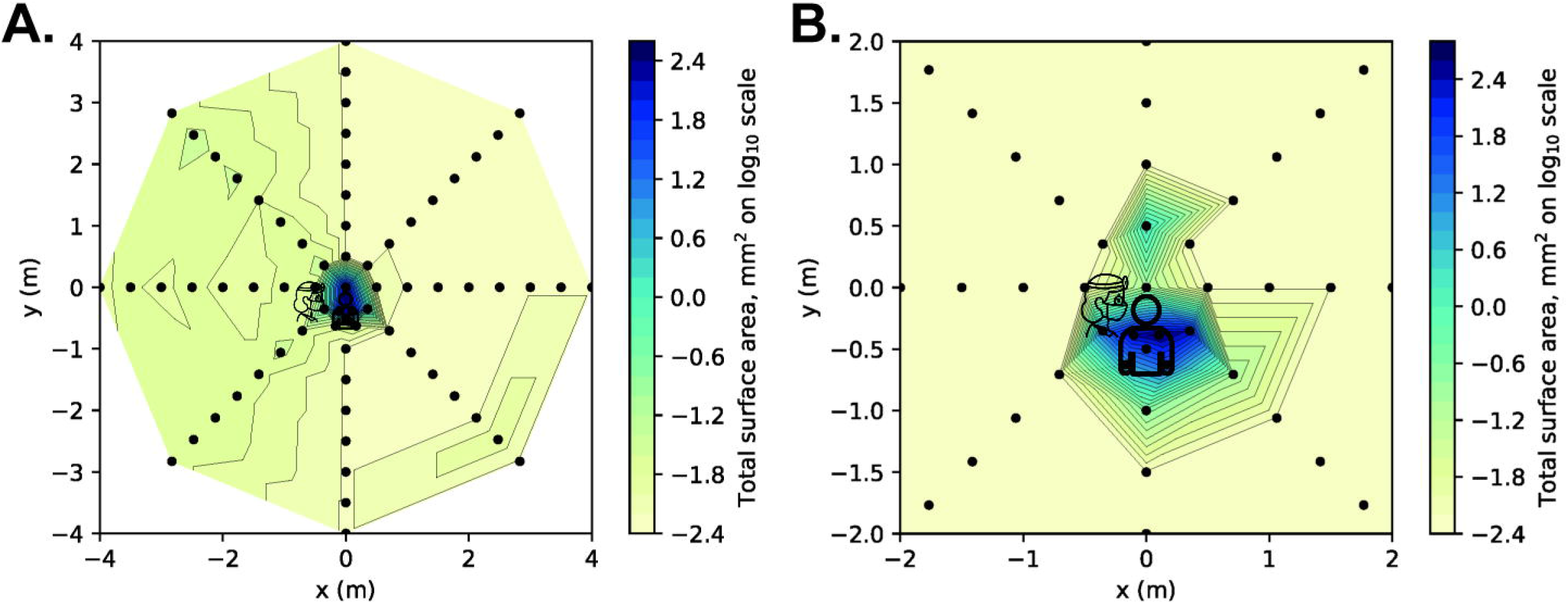
Heatmap showing surface area outcome measure for two clinical procedures. A, ultrasonic scaling. B, 3-in-1 spray. For each coordinate, the maximum value recorded from three repetitions of each clinical procedure was used as this was deemed most clinically relevant. Logarithmic transformation was performed on the data (Log_10_). Note the scale is reduced to remove areas showing zero readings in panel B only.

**Table 1.**
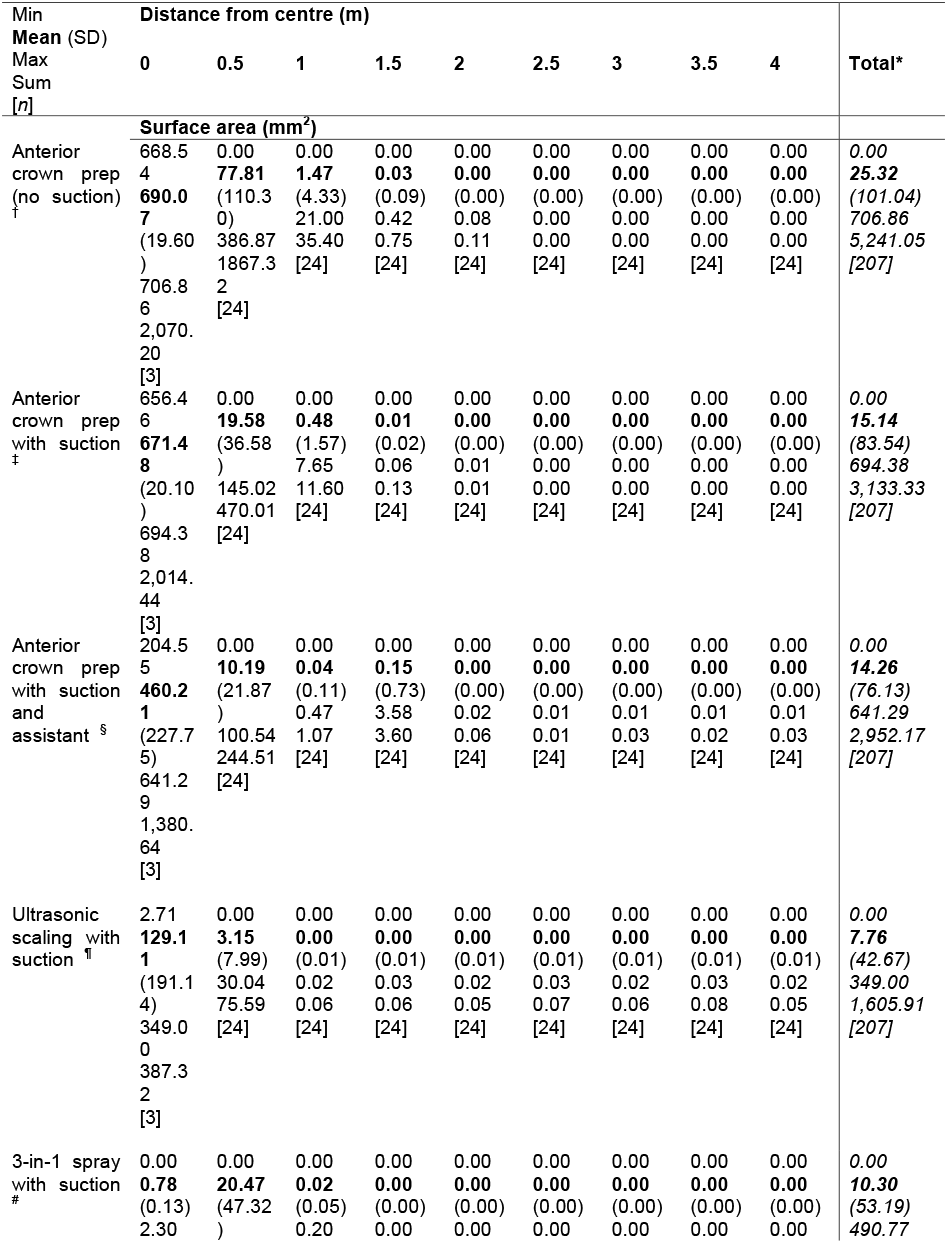
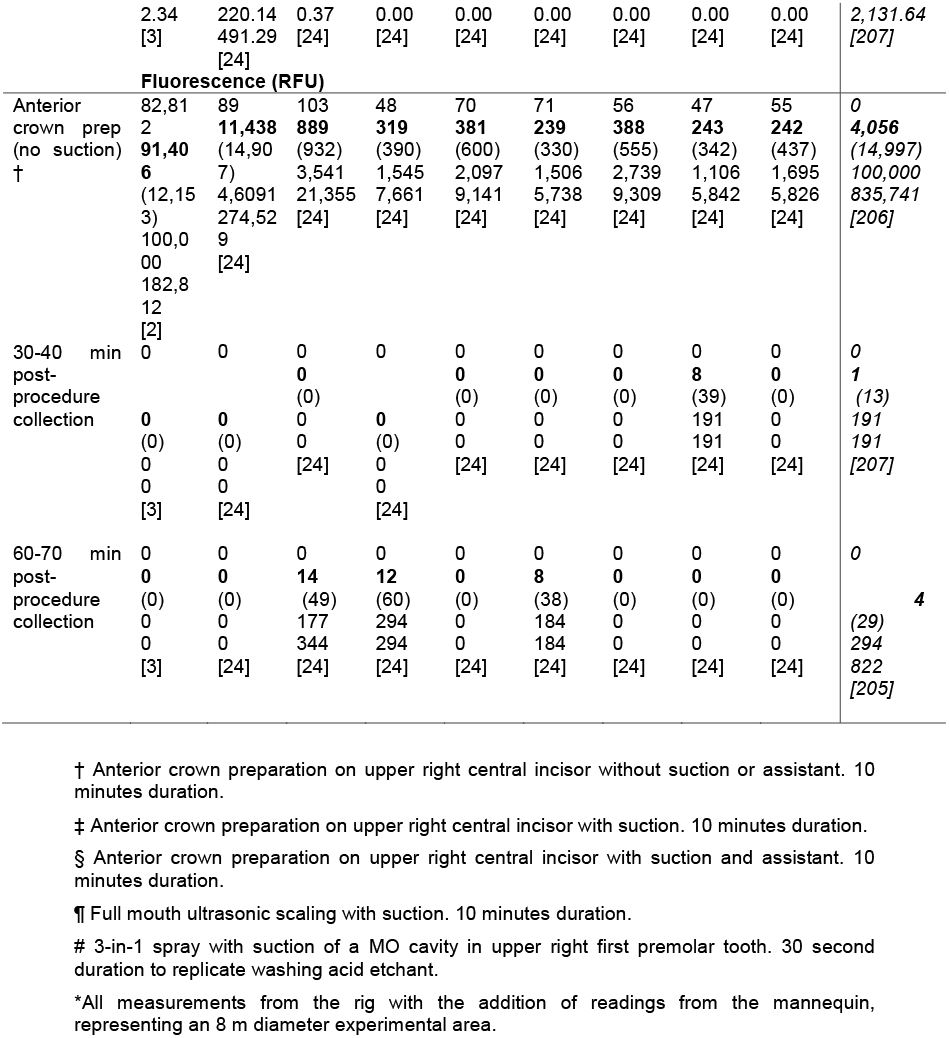
Dental aerosol and splatter as measured by contaminated surface area using image analysis or by spectrofluorometric analysis. For each experimental condition, the data from an average of three repetitions for all samples at each distance are included together. A total is also given for all samples for each condition, which also includes data from samples placed on the mannequin.

For one experimental condition (anterior crown prep with **<u>no</u>** suction, representing a presumed worst-case scenario), at three time points, we also completed spectrofluorimetric analysis (table 1). The particle count was weakly correlated with spectrofluorimetric measurements (*r*=0.344, *n*=244, *p*<0.001), average particle size was moderately correlated (*r*=0.555, *n*=244, *p*<0.001) and total surface area was very strongly correlated (*r*=0.930, *n*=244, *p*<0.001), supporting our use of surface area as the main outcome measure from image analysis (supplementary figure S1). Data from one time point are presented in Figure 4. Using serial dilution of fluorescein, we derived a standard curve covering the range 50 nM to 102 μM. The equation *y* = 700.42 *x* – 1449.5 was derived from the standard curve (*y*=fluorescence, RFU; *x*=fluorescein concentration, μM). For illustrative purposes we detail these for the 270 degree axis (mean values across three repetitions from the initial time point): 0 m = 132.6 μM; 0.5 m = 26.3 μM; 1 m = 5.25 μM; 1.5 m = 3.02 μM; 2 m = 3.44 μM; 2.5 m = 3.30 μM; 3 m = 2.79 μM; 3.5 m = 2.86 μM; 4 m = 3.09 μM.

**Figure 4.**
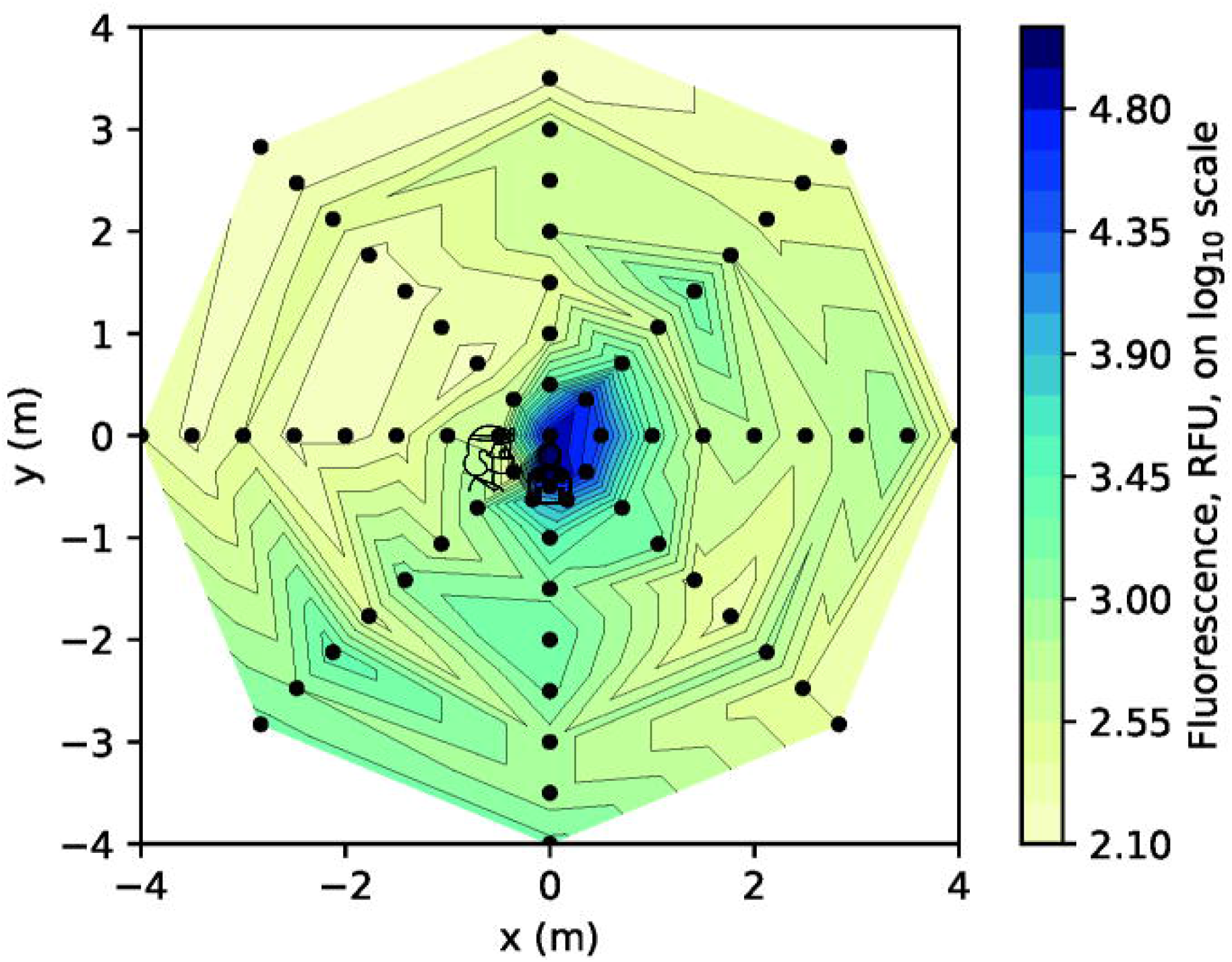
Heatmap presenting spectrofluorimetric analysis of the samples from the anterior crown preparation (without suction) clinical procedure at 0 – 10 minutes (surface area data shown in Figure 2A). For each coordinate, the maximum value recorded from three repetitions of each clinical procedure was used. Logarithmic transformation was performed on the data (Log_10_). Note the scale includes the full dimensions of the experimental rig. RFU: relative fluorescence units.

The mannequin, operator and assistant were all heavily contaminated (supplementary table S2). The operator’s left (non-dominant) arm, left body and lower visor were the most contaminated sites. Generally, levels of contamination were much lower for the assistant, being highest on the left arm and left chest (the assistant used their right hand to hold the suction tip, and left hand to support the tubing). All areas of the mannequin were heavily contaminated. The operator and assistant’s masks (only assessed in one condition) showed low but measurable contamination, usually at the lateral edges.

### Effect of dental suction (with and without assistant)

The use of dental suction, held by the operator, reduced the contamination of filter papers at each distance (table 1), although image analysis still detected contamination up to 2 m. Between 0.5-1.5 m there was a 67-75% reduction (central site contamination was unaffected). The spatial distribution was altered as demonstrated in figure 2. When an assistant was present and held the dental suction this further reduced contamination readings within the first 1m, however, we noted a marked increase at the 1.5 m reading behind the assistant (0°).

### Procedure type

Three clinical procedures (anterior crown preparation, ultrasonic scaling, and 3-in-1 spray use) were assessed while the operator held dental suction. The highest readings were obtained from the anterior crown preparation, but each procedure gave a unique distribution (tables 1 and S2; figures 2 and 3). The ultrasonic scaler produced high levels of contamination at the centre, reducing markedly at 0.5 m, but with low levels of contamination detectable up to the 4 m limit of measurement. The 3-in-1 spray procedure produced high levels of contamination at 0.5 m but little beyond 1 m.

### Effect of time

Image analysis demonstrated no detectable fluorescein contamination of the filter papers at 30-40 and 60-70 minutes post procedure (for the anterior crown preparation without suction condition). Additionally, spectrofluorometric analysis of these samples demonstrated very low levels of contamination. The overall contamination across the 8 m diameter experimental area at 30-40 minutes was 0.02% of the original level, and at 60-70 minutes it was 0.10% of the original level (Table 1).

### Particle size

Average particle size measurements were combined for the 0, 0.5, 1 and 1.5 m readings for each condition to give an indication of the nature of the particles in this area. The anterior crown preparation without suction produced the largest particles (mean ± SD: 0.49 ± 2.98 mm^2^) which were similar to when suction was added by the operator (0.56 ± 3.34 mm^2^). There was a size reduction when an assistant provided suction (0.11 ± 0.69 mm^2^). The ultrasonic scaling produced the smallest particles (0.05 ± 0.24 mm^2^) followed by the 3-in-1 spray (0.08 ± 0.25 mm^2^). Figure 5 presents images of all samples for one repetition of a single experimental condition to demonstrate the distribution of particles and size.

**Figure 5.**
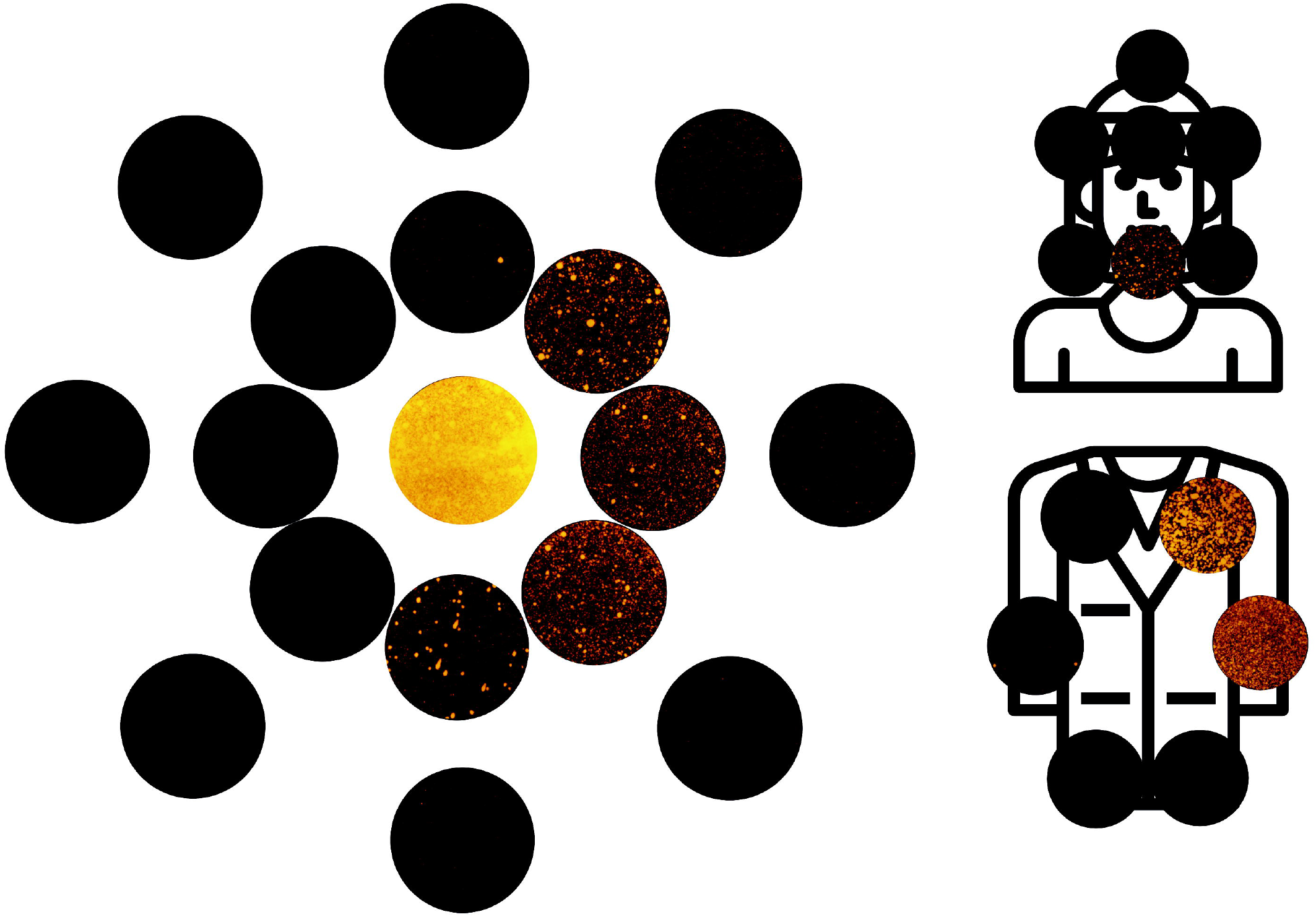
Composite image comprised of all filter paper samples images used for image analysis within the first 1 m from the one repetition of the anterior crown preparation (no suction) condition. Colour balance and contrast adjusted to aid visualisation. Samples are arranged with the central sample in the centre, and samples from 0.5 m and 1 m arranged concentrically moving outwards. The axis is the same as demonstrated in figures 1-4.

## Discussion

Dental aerosol and splatter are an important potential mode of transmission for many pathogens, including SARS-CoV-2. Understanding the risk these phenomena pose is vitally important in the reintroduction of dental services in the current COVID-19 pandemic. Our study is novel in that we are the first to measure aerosol and splatter distribution at distances up to 4 m from the source, and the first to apply image and spectrofluorometric analysis to the study of dental aerosol and splatter. This has allowed us to gather urgently needed data relevant to the provision of dental services during the COVID-19 pandemic, and more widely. Specifically, we have demonstrated the relative distribution of aerosol and splatter following different dental procedures, the effect of suction and assistant presence, and the persistence of aerosol and splatter over time.

Previous investigators have used various tracer dyes and visual examination techniques to evaluate ‘dental aerosol’ and have demonstrated positive readings at up to 1.2 m^41, 43^. Our study further optimises these methods and we have demonstrated positive readings at up to 2 m (and low levels at up to 4 m in the case of ultrasonic scaling). This is consistent with the findings of other investigators using bacterial culture methods to detect contamination at up to 2 m^36, 37, 50^. Importantly, our spectrofluorometric analysis demonstrates that some fluorescein contamination may occur beyond this on filter papers that appear clean by image analysis. We propose that studies which use dye tracers assessed by visual examination or image analysis techniques are assessing primarily splatter (particles >5 μm) rather than aerosol (<5 μm); this is because in order for deposits to be visible to the eye or camera it has to be relatively large in size (i.e. splatter). Previous research using these methods should therefore be interpreted in this context. It is, however, worth noting that larger particles are likely to contain a greater viral load, and given the risk of SARS-CoV-2 transmission through contact with mucosal surfaces^51^, from a cross infection perspective splatter is likely to be highly significant. Reassuringly, in our study splatter was greatly reduced using of suction.

Findings from both analytical techniques demonstrate contamination at a distance from the source although contamination was lower at greater distances; this shows the potential for pathogens to travel a similar distance, although our methods replicate a worst-case scenario. Within closed surgery environments this reinforces the need for minimal clutter and strict cross-infection control measures. Within open clinic environments further research is required to investigate parameters such as the impact of partitions on aerosol and splatter.

We demonstrated significant contamination of the operator, assistant and mannequin for all procedures, which is consistent with the findings of other investigators^33, 37, 40, 43^. This is unsurprising and underscores the need for adequate personal protective equipment (PPE), for the operator and assistant. This also highlights the importance of enhanced PPE^52^ during the peak of a pandemic for AGPs, because of the likelihood of treating an asymptomatic carriers. Coverage of the operator and assistant’s exposed arms with a waterproof covering would protect against contamination, although scrupulous hygiene with an effective antiseptic (povidone-iodine or 70% alcohol^53, 54^) would be a minimum requirement if this were not used. PPE for patients’ clothes do not feature in dental guidelines relating to COVID-19, and our findings would suggest significant contamination of the patient is likely during AGPs, presenting a risk of onward cross-contamination by contact with surroundings; it is therefore important to provide waterproof protection for patients’ clothes.

Our findings demonstrate that use of a high-speed air-turbine, ultrasonic scaler and 3-in-1 spray are all AGPs. 3-in-1 use is not currently included in the list of defined healthcare related AGPs recently updated by Health Protection Scotland^30^, which only details “high speed devices such as ultrasonic scalers and high-speed drills”. The highest levels of contamination were from the air-rotor, although the ultrasonic scaler demonstrated contamination at further distances, in keeping with the findings of Bennett *et al.*^20^. Dental suction was effective at reducing fluorescein contamination, with reduction of 67-75% between 0.5-1.5m. This is consistent with the effect of suction demonstrated by other investigators^36, 55^.

When dental suction was provided by an assistant this was more effective in reducing contamination, although increased readings were seen at 1.5 m, potentially indicating that an additional barrier in the form of an assistant may have a more complex aerodynamic effect. High-volume dental suction is recommended in most dental guidelines and SOPs relating to COVID-19, as an essential mitigation procedure when conducting AGPs.

However, we are not aware of any that provide a definition or basic minimal requirements for effective high-volume dental suction. National guidelines^56^ classify suction systems based on air flow rate (high-volume systems: 250 L/min at the widest bore size of the operating hose). We did not have a suitable device available to measure air flow rate of the system used in the present study and hence we chose to use the term ‘dental suction’ as we were unable to confirm whether it met this definition. We did, however measure water flow rate (6.3 L/min) which we found to be similar to that reported by other investigators^38^. Our findings highlight the importance of suction as a mitigation factor in splatter and aerosol distribution following dental procedures, and future research should examine the impact of this effect in relation to different levels of suction based on air flow rate.

Safe times following procedures, after which contamination becomes negligible have rarely been investigated robustly. In studies using tracer dyes we are only aware of a single paper reporting contamination at 30 minutes^43^. This conflicts with our findings of **<u>no</u>** contamination by image analysis at 30 and 60 minutes, and only very low levels by spectrofluorometric analysis (≤ 0.10% of original levels). It is unclear from the methods of Veena *et al.*^43^ whether new filter papers were placed immediately following the procedure and collected at 30 minutes, or placed at 30 minutes and collected thereafter; in the prior case, any contamination found on the samples could have arisen at any time from the end of the procedure up to 30 minutes, and it cannot therefore be determined when contamination actually occurred. In addition, the authors do not report whether the tape they used to support filter papers was replaced following the initial exposure, and if not, it is possible that existing contamination was transferred to filter papers placed subsequently. Finally, the investigation reported by Veena *et al.*^43^ was a single experiment and did not use multiple repetitions. It is important to note that our findings relate to the environmental setting studied, with 6.5 air changes per hour. Air exchange rates in dental surgeries are likely to vary which may affect translation.

Our study has several limitations and our results need to be interpreted in the context of these. Our methods serve as a model for aerosol and splatter contamination, and further work is required to confirm their biological validity. As our knowledge of the infective dose of SARS-CoV-2 required to cause COVID-19 develops, the clinical relevance of our findings need to be put into context; our understanding of this is still too basic to be able to draw definitive conclusions as to the risks posed by dental aerosol and splatter. Our experimental set up incorporated the tracer dye within the irrigation system of the dental units and represents a **<u>worst-case</u>** scenario for distribution of biological material.

In reality, a small amount of blood and saliva will mix with large volume of water irrigant creating aerosol and splatter with diluted pathogen concentration compared to blood or saliva, and a likely reduced infective potential^19^. It has been estimated that over a 15-minute exposure during dental treatment with high-speed instruments, an operator may be exposed to 0.014 – 0.12 μL of saliva^20^. Early data suggest a median SARS-CoV-2 viral load of 3.3 x 10^6^ copies per mL in the saliva of infected patients^22, 23^; taken together, this suggests that an operator **<u>without</u>** PPE at around 0.5 m from the source may be exposed to an estimated 46 – 396 viral copies during a 15 minute procedure. These data were collected from hospital inpatients, and recent data suggest that asymptomatic carriers may have lower salivary viral loads^27, 28^; similarly the average concentration of fluorescein detected by spectrofluorometric analysis past 2 m in the present study was almost two orders of magnitude lower than at 0.5 m, and so at distances beyond 0.5 m this risk is likely to be lower. Importantly, we still do not yet know what the infective dose of SARS-CoV-2 required to cause COVID-19 is.

## Conclusions

Within the limitation of this study, dental aerosol and splatter has the potential to be a cross infection risk even at a distance from the source. The high-speed air-turbine generated the most aerosol and splatter, even with assistant-held suction. Our findings suggest that it may be safe to reduce fallow times between dental AGPs in settings with 6.5 air changes per hour to 30 minutes. Future research should evaluate further procedures, mitigation strategies, time periods and aim to assess the biological relevance of this model.

## Acknowledgements

Charlotte Currie is funded by a National Institute for Health Research (NIHR) Doctoral Research Fellowship. David Edwards is funded by a NIHR Academic Clinical Fellowship. Jamie Coulter is funded by a NIHR BRC Doctoral Research Fellowship Richard Holliday is funded by a NIHR Clinical Lectureship. The views expressed are those of the authors and not necessarily those of the NHS, the NIHR or the Department of Health and Social Care. Nadia Rostami has been funded during this period by the Dunhill Medical Trust (RPGF1810/101) who kindly extended her funding to support this urgent COVID-19 related research. We would like to thank Ashling Dolan for advice on displaying the heat maps, Colm Kelleher (Department of Molecular and Cellular Biology, Harvard University, USA) for advice and expertise on Python programming, and to the engineers and tool curators at Newcastle Dental Hospital for their assistance with some practical aspects of this project. Consumables for this project were supported by internal funding from the School of Dental Sciences, Newcastle University.

## Data accessibility statement

Data available from the authors on reasonable request.

## Author Contributions

J. R. Allison, C. C. Currie, D. Edwards, J. Durham, C. J. Nile, N. Jakubovics, and R. Holliday, contributed to the conception and design of the study. J. R. Allison, C. C. Currie, D. Edwards, C. Bowes, J. Coulter, K. Pickering, E. Kozhevnikova, C. J. Nile, N. Jakubovics, N. Rostami, and R. Holliday contributed to the acquisition, analysis and interpretation of data. All authors were involved in drafting and critically revising the manuscript and have given final approval for publication. All authors agree to be accountable for all aspects of the work.

## Declarations

The authors have nothing to declare.

