## Supplementary material for "Evaluating aerosol and splatter following dental procedures: addressing new challenges for oral healthcare and rehabilitation": S1

### Supplementary Data S1

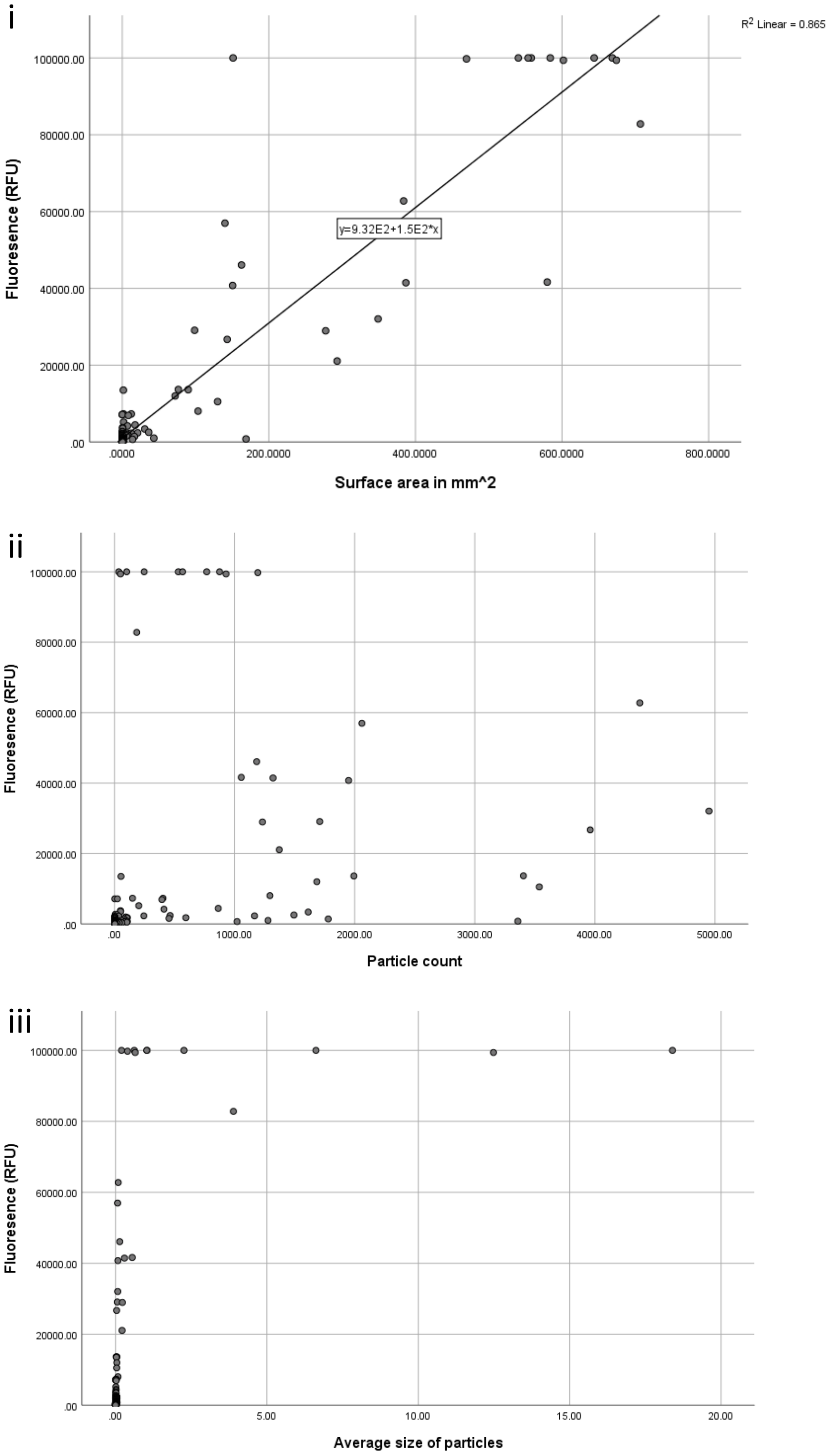


Figure S1. Correlation of image analysis data with spectrofluorometric analysis data. (i) Total surface area very strongly correlated ($r$ = 0.930, $n$ = 244, $p$ < 0.001), linear line of best fit shown; (ii) particle count weakly correlated ($r$ = 0.344, $n$ = 244, $p$ < 0.001); (iii) Particle size moderately correlated ($r$ = 0.555, $n$ = 244, $p$ < 0.001). RFU: relative fluorescence units.
