## Supplementary material for "Evaluating aerosol and splatter following dental procedures: addressing new challenges for oral healthcare and rehabilitation": S2

**Supplementary Data S2**

| | Min<br>**Mean** (SD)<br>Max<br>Sum<br>[*n*] | **Total surface area ($mm^2$)** | | | | | **Fluorescence (RFU)** | | |
|---|---|---|---|---|---|---|---|---|---|
| | | | | | | | Anterior crown prep (no suction) † | | |
| | Position on operator/assistant/ mannequin | Anterior crown prep (no suction) † | Anterior crown prep with suction ‡ | Anterior crown prep with suction and assistant § | Ultrasonic scaling with suction ¶ | 3-in-1 spray with suction # | Initial collection | 30-40 min post-procedure collection | 60-70 min post-procedure collection |
| **Mannequin** | Upper Left | 98.69 | 105.02 | 370.96 | 213.92 | 247.77 | 777 | 0 | 0 |
| | | **269.27** | **162.92** | **398.40** | **269.82** | **371.86** | **43,286** | **0** | **0** |
| | | (237.35) | (61.11) | (44.60) | (49.29) | (121.58) | (51,113) | (0) | (0) |
| | | 540.33 | 226.80 | 449.86 | 307.01 | 490.78 | 100,000 | 0 | 0 |
| | | 807.81 | 488.77 | 1,195.19 | 809.46 | 1,115.59 | 129,858 | 0 | 0 |
| | | [3] | [3] | [3] | [3] | [3] | [3] | [3] | [2] |
| | Upper Right | 43.06 | 30.40 | 19.28 | 88.91 | 41.35 | 997 | 0 | 0 |
| | | **111.44** | **35.59** | **28.71** | **111.02** | **148.61** | **52,658** | **0** | **0** |
| | | (59.47) | (8.94) | (10.68) | (35.16) | (123.87) | (49,643) | (0) | (0) |
| | | 151.13 | 45.91 | 40.30 | 151.56 | 284.91 | 100,000 | 0 | 0 |
| | | 334.31 | 106.76 | 86.14 | 333.06 | 445.83 | 157,973 | 0 | 0 |
| | | [3] | [3] | [3] | [3] | [3] | [3] | [3] | [2] |
| | Lower Left | 14.07 | 4.48 | 10.46 | 0.00 | 11.24 | 680 | 0 | 0 |
| | | **38.97** | **9.77** | **12.68** | **0.01** | **21.53** | **5,353** | **0** | **0** |
| | | (29.90) | (4.61) | (2.43) | (0.02) | (11.32) | (5,913) | (0) | (0) |
| | | 72.13 | 12.97 | 15.27 | 0.03 | 33.66 | 12,000 | 0 | 0 |
| | | 116.91 | 29.30 | 38.03 | 0.03 | 64.59 | 16,058 | 0 | 0 |
| | | [3] | [3] | [3] | [3] | [3] | [3] | [3] | [3] |
| | Lower Right | 0.82 | 1.69 | 0.02 | 0.00 | 1.38 | 743 | 0 | 0 |
| | | **2.75** | **4.11** | **0.95** | **0.01** | **3.87** | **3,213** | **0** | **0** |
| | | (2.65) | (2.45) | (0.96) | (0.01) | (4.08) | (3,572) | (0) | (0) |
| | | 5.77 | 6.59 | 1.94 | 0.02 | 8.58 | 7309 | 0 | 0 |
| | | 8.25 | 12.34 | 2.85 | 0.02 | 11.62 | 9639 | 0 | 0 |
| | | [3] | [3] | [3] | [3] | [3] | [3] | [3] | [3] |
| **Operator** | Head | 0.00 | 0.08 | 0.00 | 0.00 | 0.00 | | | |
| | | **0.14** | **0.53** | **0.20** | **0.01** | **0.00** | | | |
| | | (0.19) | (0.61) | (0.34) | (0.01) | (0.00) | | | |
| | | 0.27 | 1.23 | 0.60 | 0.02 | 0.00 | | | |
| | | 0.27 | 1.58 | 0.60 | 0.02 | 0.00 | | | |
| | | [2] | [3] | [3] | [3] | [3] | | | |
| | Left chest | 469.58 | 269.52 | 222.95 | 206.68 | 19.51 | | | |
| | | **535.70** | **484.97** | **423.03** | **308.71** | **76.51** | | | |
| | | (59.23) | (199.98) | (212.60) | (89.07) | (50.68) | | | |
| | | 583.91 | 664.65 | 646.27 | 370.95 | 116.46 | | | |
| | | 1,607.11 | 1,454.90 | 1,269.10 | 926.12 | 229.53 | | | |
| | | [3] | [3] | [3] | [3] | [3] | | | |
| | Right chest | 0.53 | 0.00 | 0.00 | 1.38 | 0.01 | | | |
| | | **2.83** | **2.02** | **0.50** | **11.25** | **7.04** | | | |
| | | (3.71) | (3.40) | (0.85) | (8.74) | (6.42) | | | |
| | | 7.12 | 5.95 | 1.49 | 18.01 | 12.58 | | | |
| | | 8.50 | 6.07 | 1.50 | 33.75 | 6.34 | | | |
| | | [3] | [3] | [3] | [3] | [3] | | | |

### Evaluating aerosol and splatter following dental procedures: addressing new challenges for oral healthcare and rehabilitation

Allison JR, Currie CC, Edwards DC, Bowes C, Coulter J, Pickering K, Kozhevnikova E, Durham J, Nile CJ, Jakubovics N, Rostami N, Holliday R

**Supplementary Data S2**

| | | | | | |
|---|---|---|---|---|---|
| Left arm | 558.02 | 638.73 | 352.45 | 9.05 | 1.74 |
| | **638.17** | **660.86** | **527.30** | **17.02** | **9.04** |
| | (77.48) | (27.20) | (157.84) | (8.46) | (7.87) |
| | 712.67 | 691.23 | 659.26 | 25.89 | 17.37 |
| | 1,875.91 | 1,982.57 | 1,581.91 | 51.07 | 27.11 |
| | [3] | [3] | [3] | [3] | [3] |
| Right arm | 0.00 | 0.79 | 0.01 | 2.35 | 0.01 |
| | **9.59** | **1.05** | **0.60** | **11.25** | **7.04** |
| | (8.57) | (0.40) | (1.01) | (13.47) | (6.42) |
| | 16.51 | 1.51 | 1.76 | 26.75 | 12.58 |
| | 28.78 | 3.15 | 1.79 | 33.74 | 21.12 |
| | [3] | [3] | [3] | [3] | [3] |
| Left leg | 0.00 | 0.00 | 0.00 | 40.44 | 77.98 |
| | **0.03** | **1.28** | **30.92** | **194.70** | **173.71** |
| | (334.69) | (2.09) | (46.90) | (208.73) | (118.20) |
| | 579.72 | 3.70 | 84.89 | 432.20 | 305.83 |
| | 579.75 | 3.85 | 92.75 | 584.09 | 521.12 |
| | [3] | [3] | [3] | [3] | [3] |
| Right leg | 0.00 | 0.00 | 0.00 | 0.00 | 0.00 |
| | **0.76** | **0.02** | **0.01** | **0.14** | **1.06** |
| | (0.31) | (0.03) | (0.02) | (0.02) | (1.00) |
| | 2.27 | 0.05 | 0.03 | 0.04 | 2.00 |
| | 2.27 | 0.05 | 0.03 | 0.04 | 3.19 |
| | [3] | [3] | [3] | [3] | [3] |
| Visor- upper | 0.00 | 0.00 | 0.00 | 0.00 | 0.00 |
| right | **0.02** | **0.01** | **0.03** | **0.01** | **0.00** |
| | (0.03) | (0.02) | (0.05) | (0.02) | (0.00) |
| | 0.05 | 0.03 | 0.09 | 0.03 | 0.00 |
| | 0.07 | 0.03 | 0.09 | 0.03 | 0.00 |
| | [3] | [3] | [3] | [3] | [3] |
| Visor-upper mid | 0.00 | 0.00 | 0.00 | 0.00 | 0.00 |
| | **0.19** | **0.10** | **0.01** | **0.01** | **0.00** |
| | (0.62) | (0.17) | (0.02) | (0.01) | (0.00) |
| | 1.16 | 0.30 | 0.03 | 0.02 | 0.00 |
| | 1.35 | 0.30 | 0.03 | 0.02 | 0.00 |
| | [3] | [3] | [3] | [3] | [3] |
| Visor- upper left | 0.00 | 0.00 | 0.00 | 0.00 | 0.00 |
| | **0.79** | **0.01** | **0.01** | **0.01** | **0.00** |
| | (0.96) | (0.02) | (0.01) | (0.01) | (0.00) |
| | 1.86 | 0.03 | 0.02 | 0.03 | 0.00 |
| | 2.36 | 0.03 | 0.03 | 0.03 | 0.00 |
| | [3] | [3] | [3] | [3] | [3] |
| Visor- lower left | 8.33 | 0.09 | 0.12 | 0.00 | 0.00 |
| | **12.74** (4.63) | **8.02** | **1.84** | **0.83** | **0.01** |
| | 17.56 | (6.86) | (2.89) | (1.02) | (0.01) |
| | 38.22 | 12.00 | 5.18 | 1.97 | 0.02 |
| | [3] | 24.05 | 5.52 | 2.49 | 0.02 |
| | | [3] | [3] | [3] | [3] |
| Visor- lower | 348.98 | 17.92 | 11.41 | 74.03 | 0.00 |
| mid | **444.87** | **63.61** | **70.51** | **96.06** | **12.43** |
| | (137.03) | (47.68) | (101.58) | (33.92) | (19.89) |
| | 601.81 | 113.05 | 187.78 | 135.12 | 35.37 |
| | 1,334.60 | 190.82 | 211.53 | 288.19 | 37.30 |

### Evaluating aerosol and splatter following dental procedures: addressing new challenges for oral healthcare and rehabilitation

Allison JR, Currie CC, Edwards DC, Bowes C, Coulter J, Pickering K, Kozhevnikova E, Durham J, Nile CJ, Jakubovics N, Rostami N, Holliday R

**Supplementary Data S2**

| | | | | | | |
|---|---|---|---|---|---|---|
| | | [3] | [3] | [3] | [3] | [3] |
| | Visor- lower right | 0.01<br>**0.15**<br>(0.23)<br>0.42<br>0.45<br>[3] | 0.00<br>**0.01**<br>(0.02)<br>0.03<br>0.03<br>[3] | 0.00<br>**0.04**<br>(0.04)<br>0.09<br>0.12<br>[3] | 0.00<br>**0.01**<br>(0.01)<br>0.02<br>0.02<br>[3] | 0.00<br>**0.00**<br>(0.00)<br>0.00<br>0.00<br>[3] |
| | Mask- left | | | 0.00<br>**0.03**<br>(0.03)<br>0.06<br>0.10<br>[3] | | |
| | Mask- mid | | | 0.00<br>**0.00**<br>(0.01)<br>0.01<br>0.01<br>[3] | | |
| | Mask- right | | | 0.00<br>**0.00**<br>(0.01)<br>0.01<br>0.01<br>[3] | | |
| **Assistant** | Head | | | 0.00<br>**0.01**<br>(0.01)<br>0.02<br>0.02<br>[3] | | |
| | Left chest | | | 0.12<br>**21.94**<br>(37.70)<br>65.46<br>65.81<br>[3] | | |
| | Right chest | | | 0.04<br>**0.14**<br>(0.14)<br>0.30<br>0.42<br>[3] | | |
| | Left arm | | | 0.00<br>**11.52**<br>(15.37)<br>28.97<br>34.56<br>[3] | | |
| | Right arm | | | 0.10<br>**4.29**<br>(3.99) | | |

### Evaluating aerosol and splatter following dental procedures: addressing new challenges for oral healthcare and rehabilitation

Allison JR, Currie CC, Edwards DC, Bowes C, Coulter J, Pickering K, Kozhevnikova E, Durham J, Nile CJ, Jakubovics N, Rostami N, Holliday R

**Supplementary Data S2**

| | |
|---|---|
| | 8.04<br>12.87<br>[3] |
| Left leg | 0.08<br>**0.58**<br>(0.48)<br>1.04<br>1.74<br>[3] |
| Right leg | 0.00<br>**0.37**<br>(0.64)<br>1.12<br>1.12<br>[3] |
| Visor- upper right | 0.00<br>**0.00**<br>(0.00)<br>0.00<br>0.00<br>[3] |
| Visor-upper mid | 0.00<br>**0.00**<br>(0.00)<br>0.00<br>0.00<br>[3] |
| Visor- upper left | 0.00<br>**0.00**<br>(0.00)<br>0.00<br>0.00<br>[3] |
| Visor- lower left | 0.00<br>**0.00**<br>(0.00)<br>0.00<br>0.00<br>[3] |
| Visor- lower mid | 0.00<br>**0.02**<br>(0.03)<br>0.05<br>0.05<br>[3] |
| Visor- lower right | 0.00<br>**0.00**<br>(0.01)<br>0.01<br>0.01<br>[3] |
| Mask- left | 0.00 |

### Evaluating aerosol and splatter following dental procedures: addressing new challenges for oral healthcare and rehabilitation

Allison JR, Currie CC, Edwards DC, Bowes C, Coulter J, Pickering K, Kozhevnikova E, Durham J, Nile CJ, Jakubovics N, Rostami N, Holliday R

**Supplementary Data S2**

| | |
|---|---|
| | **0.01**<br>(0.01)<br>0.01<br>0.01<br>[3] |
| Mask- mid | 0.00<br>**0.00**<br>(0.00)<br>0.00<br>0.00<br>[3] |
| Mask- right | 0.00<br>**0.02**<br>(0.04)<br>0.07<br>0.07<br>[3] |

Table S2. summary of surface area and spectrofluorometric data collected from the mannequin, operator and assistant. Each location represents three repetitions. RFU: relative fluorescence units.

† Anterior crown preparation on upper right central incisor without suction or assistant. 10 minutes duration.

‡ Anterior crown preparation on upper right central incisor with suction. 10 minutes duration.

§ Anterior crown preparation on upper right central incisor with suction and assistant. 10 minutes duration.

¶ Full mouth ultrasonic scaling with suction. 10 minutes duration.

### 3-in-1 spray with suction of a MO cavity in upper right first premolar tooth. 30 second duration to replicate washing acid etchant.
